# Prevalence of mango postharvest Stem-end rot disease in Cote d’Ivoire and identification of fungal pathogens associated

**DOI:** 10.1101/2023.08.02.551668

**Authors:** Yéfoungnigui S. Yeo, Yassogui Kone, Dio D. Dembele, Elisee L D G Amari, Jean-Yves Rey, Emerson M. Del Ponte, Diana Fernandez, Daouda Kone

**Author notes:** co-last author (Diana Fernandez, Daouda Kone).

## Abstract

The Stem-end rot (SER) postharvest disease of mango (*Mangifera indica* L.) fruits is a significant economic threat to mango production. If suitable conditions are not maintained, it can lead to losses of up to 100 %. Despite its importance, very little information is known about this disease in Côte d’Ivoire. This research aimed determining the incidence and severity of SER in mango orchards, assess how preharvest climate parameters affect the disease and determine the pathogenic fungi associated with SER. Therefore, mango SER was evaluated on 1500 mango fruits collected from 15 orchards.in Boundiali, Ferkéssédougou, Korhogo, Odienné, and Sinématiali departments. Mango SER incidence ranged from 10 % to 30 %, while severity ranged from 5 % to 20 %. No significant differences in these parameters were observed between the different departments (P>0.05). The study also revealed a low correlation between SER disease incidence and mean air temperature (r=0.36) and minimum air temperature (r=0.26) data, indicating that preharvest weather conditions may have a marginal impact on mango SER disease intensity in the postharvest phase. Pathogenic fungi associated with SER were isolated and identified using morphological characteristics and multilocus sequence analysis of the rDNA internal transcribed spacer (ITS) region and the translation elongation factor 1-alpha (*tef1-α*). We identified various fungal species associated with mango SER disease, with *Lasiodiplodia* species (74%) being the most prevalent (including *Lasiodiplodia theobromae, L. euphorbicola*, and *L. caatinguensis*), followed by *Colletotrichum gloeosporioides*, *Curvularia pseudobrachyspora*, *Diaporthe endophytica* and *Fusarium mangiferae*. However, only *Lasiodiplodia* species and *Diaporthe endophytica* induced SER symptoms. This study was the first ever evaluation of mango SER disease and associated fungal pathogens identification in Côte d’Ivoire. This result will assist researchers in developing a control method for mango SER.

## 1 Introduction

Mango (*Mangifera indica* L.) is a highly valued fruit produced in tropical, subtropical, and semi-arid regions. According to FAOSTAT (2021), the global production of mangoes is estimated at over 57 million tons. Mangoes are rich in nutrients such as fibre, vitamins C and A, polyphenols, carotenoids, carbohydrates, and calcium (Owino and Ambuko, 2021). However, mango production is threatened by several fungal diseases, including postharvest Stem-end rot (SER). The SER disease is characterized by small dark-brown lesions in the peel surrounding the fruit stem end, eventually leading to soft and watery decay, causing complete fruit rot (Galsurker et al., 2020). Without proper management strategies, the disease can significantly reduce fruit quality and may result in postharvest losses of up to 100% in some orchards (Alam et al., 2017; Honger and William, 2015).

Several fungal species have been associated with mango SER, including *Neofusicoccum brasiliense*, *N. parvum*, *N. mangiferae*, *Fusicoccum aesculi*, *Nattrassia mangiferae*, *Neoscytalidium dimidiatum*, *Lasiodiplodia theobromae*, *Curvularia* sp, *Fusarium* sp, *Neocosmospora* sp, *Pestalotiopsis* sp, *Neopestalotiopsis* sp, *Colletotrichum gloeosporioides*, and *Alternaria* sp. (Adikaram et al., 2022b; Alam et al., 2017a; Phillips et al., 2013; Marques et al., 2013; Johnson et al., 1992). The pathogen may colonize inflorescence and pedicel tissues during flowering and remain as endophyte in phloem and xylem vessels (Johnson et al., 1992). Infected fruits may remain asymptomatic until ripening, making detection difficult for farmers during harvest (Govender, 2004). As a result, the disease is often only identified during storage and transportation, which is a challenge for effective control (Johnson, 2008; Terao et al., 2018).

Mango production contributes significantly to the economy of Côte d’Ivoire, generating over 10 million euros of income for local communities, corresponding to 4 % of the country’s gross domestic product (GDP) (Pugnet, 2018). The country is also the third largest supplier of mangoes to the European Union market, after Brazil and Peru (Pugnet, 2018). However, despite government and private sector efforts, postharvest losses remain a major challenge for mango production, with global losses ranging from 30 to 35 % of the national production in Côte d’Ivoire, corresponding to 5 millions euros (Kouassi, 2012). Reducing these losses through control strategies can significantly enhance the mango sector contribution to the country economy and improve the livelihoods of local communities. Prior research in Côte d’Ivoire on mango postharvest fungal diseases has mainly focused on anthracnose diseases caused by *Colletotrichum* species (Dembélé *et al*., 2020; Kouamé et al., 2011; N’Guettia & Kouassi, 2014). While anthracnose may be well controlled with fungicide application, mango SER still has a significant economic impact for producing countries (Galsurker et al., 2018; Karunanayake & Adikaram, 2020).

Several factors, including climatic parameters such as temperature and relative humidity have been reported that affect mango SER (Diedhiou et al., 2007; Alkan, 2018), probably by impacting directly pathogen infection(Riquelme-Toledo et al.,2020).. We hypothesize that weather parameters may impact the inflorescence colonization rate by endophytic fungi during flowering, which could influence the incidence and severity of SER. Understanding the potential impact of weather parameters on SER disease is essential for developing effective control strategies. The objective of this research is three-fold: firstly, to asess the incidence and severity of mango SER in Côte d’Ivoire; secondly, to examine the relationships between preharvest climate conditions and the incidence and severity of SER; and thirdly, to identify the specific fungal agents causing SER.

## 2 Material and methods

### 2.1 Orchards surveys and SER evaluation

Mango SER was evaluated on mango fruits (cv. Kent) at physiological maturity, collected from 15 mango orchards located in the departments of Korhogo, Sinématiali, Ferkéssédougou, Boundiali, and Odienné. These regions are the main mango production areas in the country. Three orchards located at least 5 km apart with a similar age (>30 years) were selected in each department. Then 100 fruits were harvested from 20 randomly selected trees in each orchard.

Fruits external surface was disinfected using a 5% hypochlorite solution, then rinsed with tap water and air dried for 30 minutes. Fruits were then packed by orchards and stored in a cold room with a temperature of 14.5 ± 0.5°C and relative humidity of 80 ± 2 % for two weeks to simulate sea-shipment transportation from Côte d’Ivoire to the European market. Then, fruits were transferred to another room with temperatures ranging from 31 ± 5°C and relative humidity of 70 ± 5%. SER symptoms were observed weekly for 28 days (incubation period). The severity index of each fruit was determined using a severity scale adapted from Alvindia and Acda (2015) ranking from 0 to 6, where 0 = no discoloration of the stem-end, 1 = discoloration limited at the stem-end, 2 = 10% discoloration of the fruit surface area initiated by stem-end rot, 3 = 11-30% discoloration of the fruit surface area initiated by stem-end rot, 4 = 31-50% discoloration of the fruit surface area initiated by stem-end rot, 5 = more than 51% discoloration of the fruit surface area initiated by stem-end rot, 6 = 100 % discoloration of fruit surface. The incidence (DI) and severity (DS) of postharvest SER disease were calculated for each orchard using the following formulas:

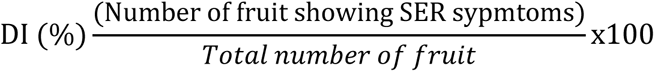

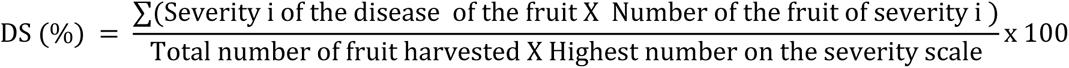

### 2.2 Climate data records

In order to evaluate the correlation between climate parameters and SER incidence and severity, we downloaded monthly climatic data such as Relative Humidity at 2 meters above ground (RH2M), Dew point temperature at 2 meters above ground (T2MDEW), maximum air temperature at 2 meters above ground (T2M_MAX), average air temperature at 2 meters above ground (T2M) and the minimum air temperature at 2 meters above ground (T2M_MIN), spanning six months prior to the survey (November to April 2021) from the National Aeronautics and Space Administration (NASA) POWER project *via* the nasapower R package version 4.0.9 (Sparks, 2018). Data were downloaded using the geographic coordinates (latitude and longitude) of the 15 orchards surveyed across the departments surveyed. The six months average data of each climatic variable were then correlated with SER postharvest incidence and severity.

### 2.3 Fungal isolation

The fruits showing SER symptoms were washed briefly using distilled water. Five fragments measuring 5×5 mm were cut from the margin between symptomatic and asymptomatic tissues. Fragments were surface sterilized by dipping them in 70 % ethanol for one minute, then in a 5 % sodium hypochlorite (NaClO) solution, and then rinsed three times with distilled sterile water. After sterilization, fragments were briefly dried on sterile filter paper. Five fragments were placed on each Petri dish (9 cm diameter) containing potato dextrose agar medium (PDA) and incubated at room temperature at 25 °C in darkness for one week. Cultures were checked daily for hyphal growth, and emerging hyphae were transferred to a new PDA plate. Fungi that did not sporulate on PDA were cultured on a 2 % water-agar (WA) containing sterilized pine (*Pinus nigra* Arnold) needles and incubated at 25 °C with a 8 h photoperiod for four weeks to induce the formation of fruiting bodies and sporulation. Fresh culture of single spore fungal pure cultures of each isolate were conserved (on toothpicks sticks at 5°C) at Félix Houphouët-Boigny University in Côte d’Ivoire.

### 2.4 Morphological identification of SER fungal isolates

The macro-morphological characteristics of fungal colonies, such as mycelium appearance, colour and spore production, were assessed from single spore-conidial cultures grown on PDA or WA. Conidia and conidiogenous cells were observed from material mounted in lactic acid under a light microscope (Zeiss Z2, Alen, Germany), and some characteristics of conidia (shape, size, colour, longitudinal striations and presence or absence of septum) were recorded. Photographs were made using an Axiocam 506 colour camera with Zeiss Image Z2 software. Morphological characters were used to identify the fungi isolated to the genus level.

### 2.5 DNA extraction, PCR amplification and sequencing

Based on morphological characters and pathogenicity test, the isolates Fr-F01; FRK37, FRZ20, FRZ27, SA9FSy, BenFFSy, and NTA were selected for molecular identification (Table 2). Fungal isolates were cultured on PDA and incubated at 25 C for 5 days. About 20 mg of mycelium was scraped and ground with a disposable pestle mounted on an electric homogenizer (Dutscher, France) into a 1.5 ml tube containing 50 µl of freshly prepared 0.5 M sodium hydroxide (NaOH). The total genomic DNA was extracted using a rapid extraction protocol of Groppe and Boller (1997). The internal transcribed spacer (ITS) and elongation translation factor (*tef1-α*) genes sequences were amplified by Polymerase chain reaction (PCR) assays and sequenced using primer pairs, ITS4 (TCCTCCGCTTATTGATATGC) / ITS5 (TCCTCCGCTTATTGATATGC) (White et al., 1990) and EF1-688F(CGGTCACTTGATCTACAAGTGC) / EF1-1251R (CCTCGAACTCACCAGTACCG) (Alves et al., 2008), respectively. For each PCR reaction, a mixture (25 µl) was prepared, which contained 5 µl of 5x Green GoTaq buffer, 2 µl of deoxynucleotide triphosphate polymerase (dNTPs) (5 mM), 1 µl of each primers (10 µM), 0.125 µl of GoTaq G2 polymerase (Promega, France), 14.875 µl of sterile Milli-Q water, and 2 µl of genomic DNA. PCR was realized with a MyCycler™ Thermal Cycler following these different steps: initial denaturation at 94 °C for 2 min, then 30 cycles of denaturation at 94 °C for 30 s (ITS) and 95 °C for 30 s (*tef1-α*), annealing at 58°C for 30 s (ITS) / 55 °C for 45 s (*tef1-α*), elongation at 72°C for 30 s (ITS) / 90 s (tef1-α), and final elongation at 72°C for 8 min (ITS)/10 min (*tef1-α*). The PCR products were separated by electrophoresis for 20 min at 100 V in a 1 % agarose gel stained with ethidium bromide. The gel was viewed and photographed using the Bio-Rad Molecular Imager® Gel Doc™ XR System and Bio-Rad Quantity One® Software. The size of the amplified PCR products was ascertained by utilizing a 100 bp GeneRulers™ DNA ladder (Promega). PCR products were sent to Genewiz/ Azenta company for DNA sequencing in both directions.

### 2.6 Sequence alignment and phylogenetic analyses

Raw ITS and *tef1*-α sequences were trimmed, aligned, and manually adjusted, and consensus sequences were generated with the BioEdit software (Hall, 1999). The genetic relatedness of the ITS and *tef1*-α of isolates sequences with other fungal sequences in the GenBank database was determined using the nucleotide Basic Local Alignment Search Tool (BLASTN) (https://blast.ncbi.nlm.nih.gov/Blast.cgi) with default parameters. All sequences obtained were deposited in the GenBank database (Table 2). Based on the best BLASTN homology results, different ITS and *tef1-α* datasets were used to assess the phylogenetic positions of SER isolates. We retrieved in GenBank the sequences from well characterized isolates (type collection) of *Lasiodiplodia*, *Colletotrichum*, *Curvularia*, *Fusarium* and *Diaporthe* species. Prior to phylogenetic analysis, ambiguous sequences at the start and end were deleted in order to optimize the alignments. Multiple sequence alignments of SER isolates and reference type collection sequences were generated using the default settings of the ClustalW algorithm in MEGA 11 software (Tamura et al., 2021). Alignment parameters were applied to find the best nucleotide substitution model in MEGA 11 with the Bayesian Information Criteria (BIC) and Akaike Information Criteria (AIC). Phylogenetic analysis was conducted using the maximum likelihood (ML) method. The Nearest Neighbor Interchange (NNI) heuristic method was used for trees inference. The initial tree was automatically generated using the Maximum Parsimony method. The ML trees were generated with 1000 bootstrap replications. All analyses were performed using MEGA 11 software (Tamura et al., 2021).

### 2.7 Pathogenicity tests

Pathogenicity tests of SER isolates were realized on mature, asymptomatic, without any visible signs of blemishes mango fruits (cv Kent). Fruits were surface sterilized by deeping them in 1 % NaClO solution for 10 min and rinsed three times with sterile distilled water before being air dried for 30 min. Using a sterile scalpel, 1 cm^2^ of surface tissue was excised from the fruit at the peduncle. A 5-mm fungal mycelial plug of 7 day-old culture of each isolate prepared on PDA was used to inoculate fruits, according to 5 fruits per isolate (with 3 replications). An agar plug without mycelium was applied on 5 fruits (with 3 repetitions) for control. The inoculated portion was wrapped with parafilm. Inoculated fruits were then incubated at 25 ± 2 °C with a relative humidity of 55 ± 2 % for 10 days. Fruits were checked daily for symptoms development. To satisfy Koch’s postulate, pieces from diseased tissues of symptomatic fruits were removed, surface sterilized as mentioned above and plated on PDA. An isolate was considered pathogenic when it was able to cause disease symptoms (rot) and further isolated from the inoculated fruit. The morphological characteristics of the isolates were compared with those of the original isolates.

### 2.8 Statistical analysis

Mango SER, incidence and severity data were checked for normal distribution and homogeneity, running Shapiro-Wilk and Levene tests, respectively. A series of analysis of variance (ANOVA) tests were performed, and in case of significant differences, means were separated using Tukey’s test (*P* <0.05). We performed the linear regression model to determine the effect of weather parameters on SER incidence and severity. Correlation analysis was used to determine the strength of the association between each pair of variables, considering all climatic variables and mango SER incidence. All analyses were performed using R software (R Core Team, 2021).

## 3 Results

### 3.1 Incidence and severity of mango SER

Mango SER disease incidence and severity were evaluated in the primary area for mango production in North West Côte d’Ivoire, covering five departments, namely Boundiali, Ferkéssédougou, Korhogo, Odienné, and Sinématiali. In each department, disease data were recorded in three different orchards (100 fruits each). SER incidence in orchards ranged from 10% to 30 %, while severity varied from 5% to 20 % (Table 1, Figure 1 A, B). The most frequent occurrence for both variables was 10 % (Figure 1 C, D). Furthermore, the analysis revealed a strong positive correlation (r = 0.92) between the SER incidence and severity (Figure 1E). Depending on departments, mango SER varied from 11% to 22 %, while severity estimates varied from 7% to 17 %. However, no significant difference was observed between departments for both variables (P > 0.05) (Table 1).

**Figure 1:**
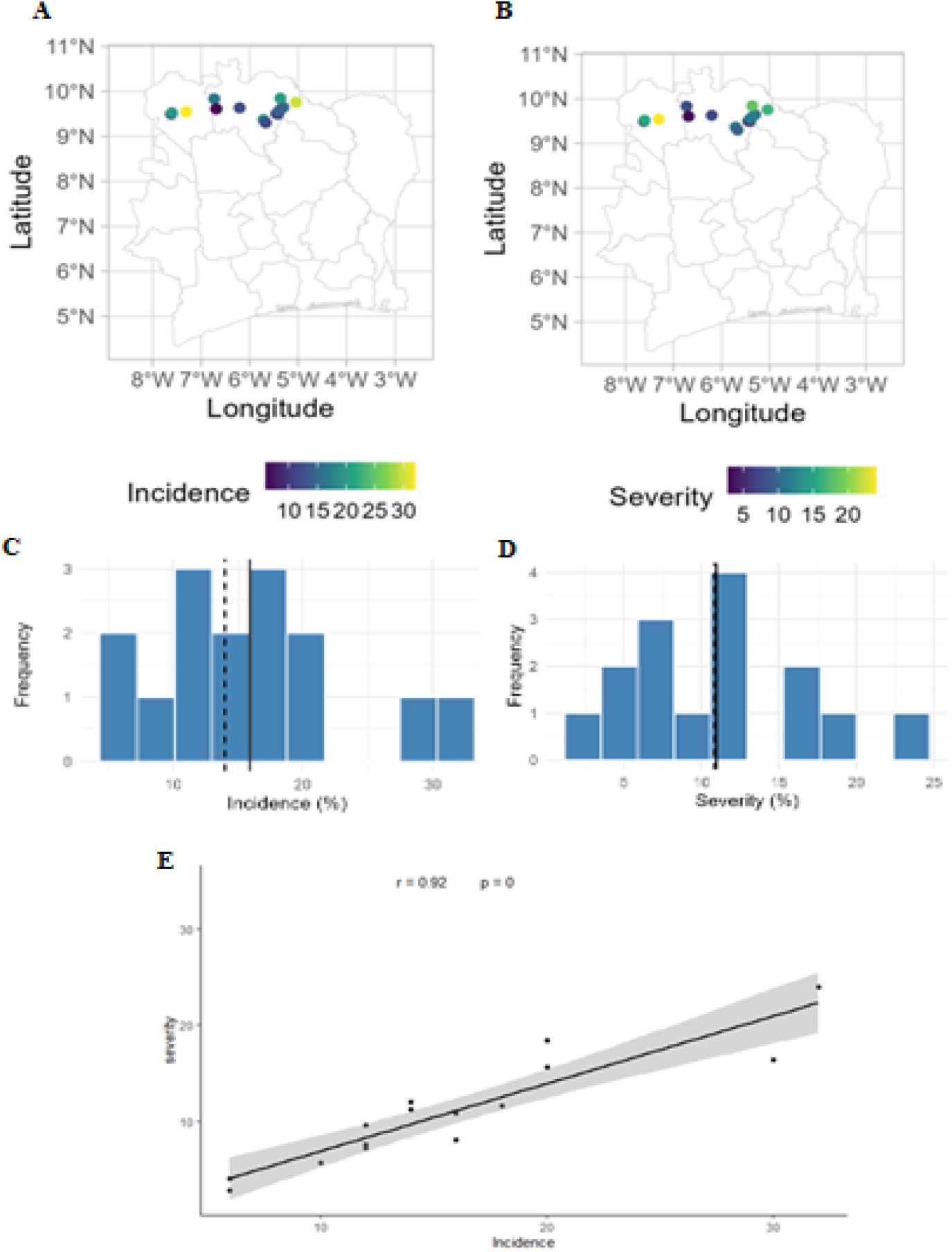
Distribution map and frequency of mango SER incidence (A, C) and severity (B, D) of mango SER postharvest disease in 15 orchards from Côte d’Ivoire and the correlation between both variables (E).

**Table 1:** Incidence and severity of SER in fifteen orchards from five departments in Côte d’Ivoire. Data presented for departments are means ± SD (n=3)

| Department | SER incidence (%) | SER severity (%) | Orchard | SER incidence (%) | SER severity (%) |
| --- | --- | --- | --- | --- | --- |
| Sinematiali | $12 \pm 2$ | $9.4 \pm 2.8$ | Klolikaha | 12 | 7.6 |
|  |  |  | Donassokaha 1 | 14 | 11.2 |
|  |  |  | Donassokaha 2 | 10 | 5.6 |
| Korhogo | $14.7 \pm 3.0$ | $11.8 \pm 1.3$ | Tioroniaradougou | 12 | 9.6 |
|  |  |  | Sissian | 18 | 11.6 |
|  |  |  | Oualokaha | 14 | 12 |
| Boundiali | $11.3 \pm 5.0$ | $7.6 \pm 2.8$ | Boundiali 1 | 12 | 7.2 |
|  |  |  | Boundiali 2 | 16 | 8 |
|  |  |  | Boundiali 3 | 6 | 2.8 |
| Odienné | $19.3 \pm 13.0$ | $19.8 \pm 10.0$ | Odienné 1 | 6 | 4 |
|  |  |  | Odienné 2 | 32 | 24 |
|  |  |  | Odienné 3 | 20 | 15.6 |
| Ferkessedougou | $22 \pm 7.2$ | $17.4 \pm 3.9$ | Village A | 16 | 10.8 |
|  |  |  | Tchékpè | 20 | 18.4 |
|  |  |  | Ferkéssédougou | 30 | 16.4 |

### 3.2 Effect of climate parameters on mango SER incidence and severity

The results of the correlation test between the average six-months climatic parameters prior to the survey and SER incidence and severity are presented in Figure 2. Our analysis revealed no correlation between PREC, RH2M, T2MDEW and T2M_MAX and the incidence and severity of SER as indicated by the null correlation coefficients (Figure 2 A, B). On the other hand, we observed a low positive correlation between T2M and T2M_MIN and the incidence of SER (Figure 2A), with correlation coefficients of 0.36 and 0.26, respectively.

**Figure 2:**
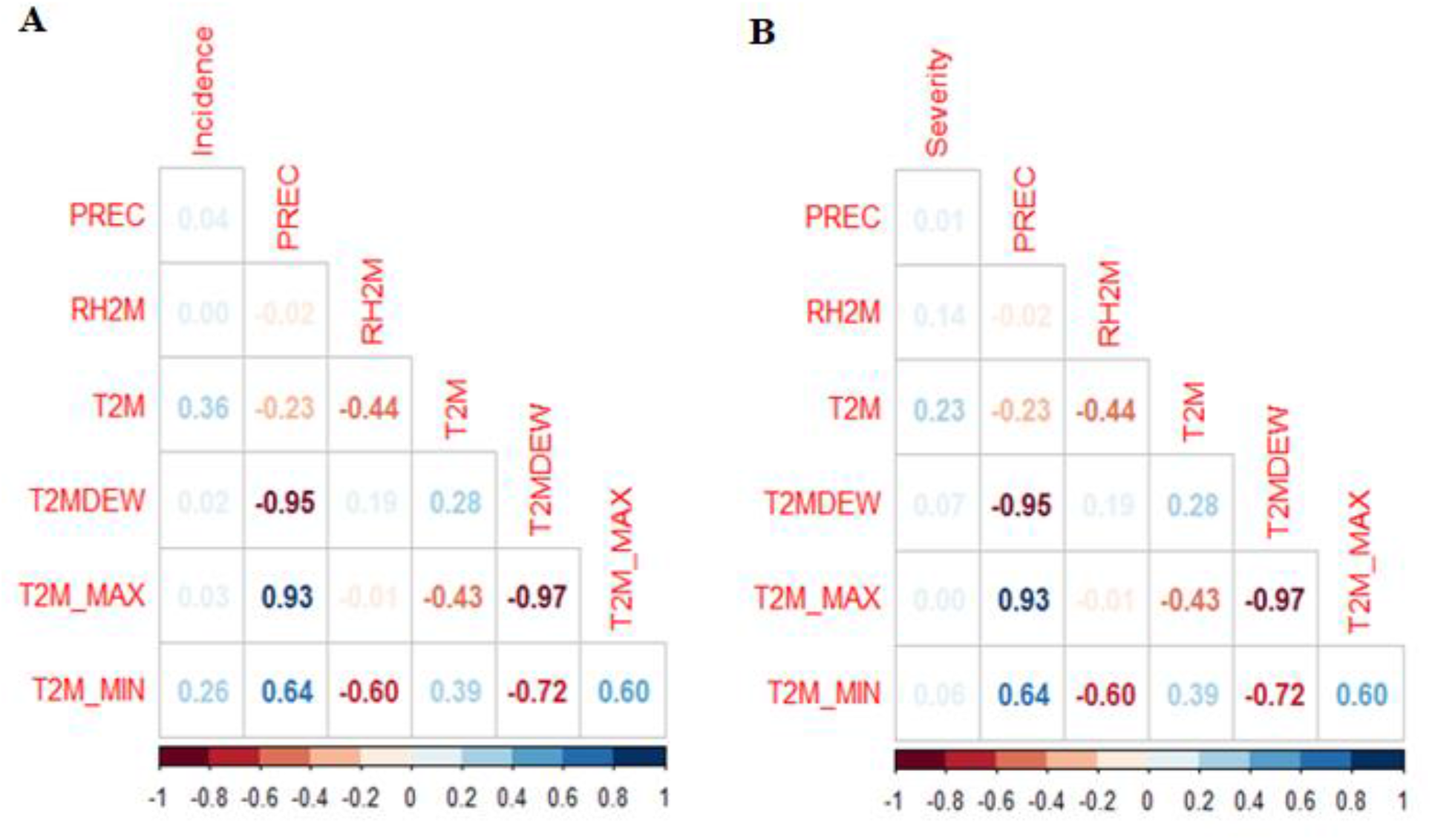
Correlation matrix of mango SER and weather parameters: A= Incidence; B= Severity. PREC= precipitation; RH2M= Relative Humidity at 2 meters above ground; T2MDEW =Dew point temperature at 2 meters above ground; T2M_MAX= maximum air temperature at 2 meters above ground; T2M= average air temperature at 2 meters above ground; T2M_MIN = minimum air temperature at 2 meters above ground.

### 3.3 Fungi identification

Fungal colonies isolated from 350 fruits presenting SER symptoms were characterized by morphological characteristics of colonies growing on PDA or pine needles on WA. Five fungal genera were identified, including *Lasiodiplodia* sp. that were the most isolated (74 %), followed by *Colletotrichum* sp. (9 %), *Curvularia* sp. (7 %), *Diaporthe* sp. (6 %) and *Fusarium* sp. (4 %).

*Lasiodiplodia* sp. colonies on PDA reached 95 mm diam in 7 days, produced dense aerial mycelium, initially white, then turned to dark brown and black (Figure 3a,b). Conidia were subovoid to ellipsoidovoid, with the apex broadly rounded, thick-walled, and contents granular; initially hyaline and aseptate, then dark brown and 1-septate with longitudinal striations (figure 3c). Conidia size was 13–30.9 μm length (avg. ±S.D = 21.8 ± 3.2 μm) × 9.0-17 μm width (avg. ±S.D = 13.0 ± 0.1 μm) (n = 50), (L/W ratio = 1.8).

**Figure 3:**
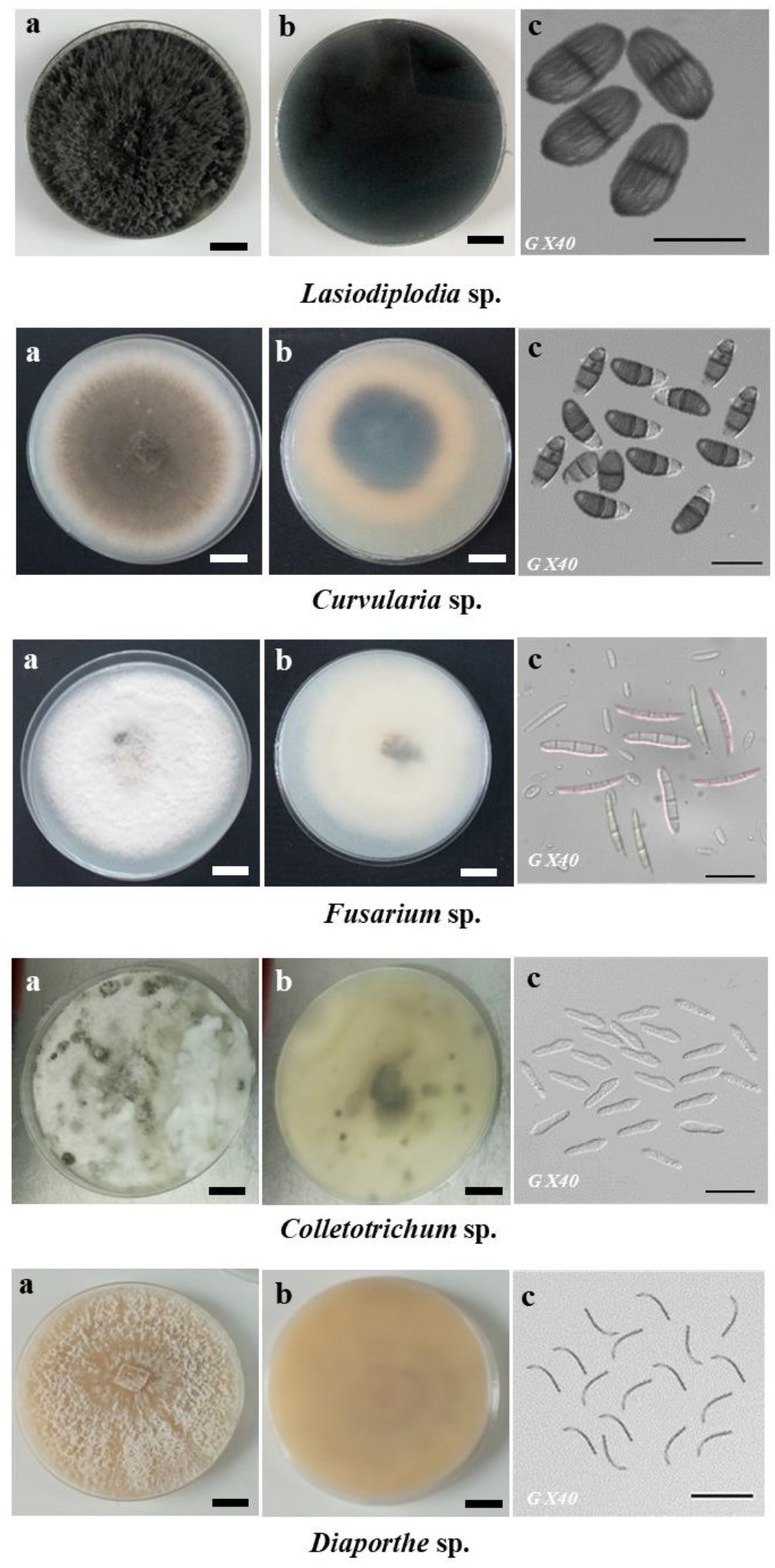
Colony and conidia morphology of *Lasiodiplodia* sp*., Curvularia* sp., *Fusarium* sp., *Colletotrichum* sp., *Diaporthe* sp. grown on PDA: (a) upper, and (b) lower surface appearance, (c) conidia. Scale bars (a, b) = 2 cm; (c) = 20 µm.

*Curvularia* sp. fungal colonies on PDA reached 65–70 mm diam in 7 days, exhibited gray, cotton-like mycelia with ring patterns on the plate (figure 8a). On the reverse plate, the pigmentation ranged from brown to blackish-brown (Figure 8b). The conidia, either straight and ovoid or fusiform in shape, had two or three septa, with the middle septum being noticeably darker (figure 3c). Conidia were 15.8–25.2 μm length (avg. ±S.D = 20.4 ± 2.3 μm) × 9-12.0 μm width (av. ±S.D = 10.5 ± 1.5 μm) (n = 50), (L/W ratio = 1.9).

*Fusarium* sp fungal colonies reached 45–80 mm diam in 7 days, displayed a white colouration (figure 3a,b). The micro and macroconidia were abundant, and their shape was hyaline, oval, or cylindrical (figure 3c). Macroconidia were 22.2-58.5 μm length (avg. ±S.D = 40.3 ± 3.2 μm) × 2-4.9 μm width (avg. ±S.D = 3.4 ± 1.1 μm) (n = 50), (L/W ratio = 11.7).

*Colletotrichum* sp. colonies on PDA reached 53–64 mm diam in 7 days, displayed a pale olivaceous colouration, ranging from grey to olivaceous grey and merging into grey olivaceous at the centre (figure 3a). The reverse of the colonies appeared as iron grey to olivaceous black (figure 3b). Conidia were unicellular, hyaline, smooth or slightly roughened. (figure 3c). Conidia were 14.8-20.5 μm length (avg. ±S.D = 17.6 ± 4.0 μm) × 5-6.2 μm width (avg. ±S.D = 5.6 ± 3.4 μm) (n = 50), (L/W ratio = 2.3).

*Diaporthe* sp. colonies reached 82–85 mm diam in 7 days, mycelium had a white-grey appearance, with small clumps of scattered pycnidia (figure 3a). The ß conidia formed were unicellular, filiform, hyaline, aseptate smooth, spindle-shaped, and slightly curved (figure 8c). Conidia were 18.7-25 μm length (avg. ±S.D = 21.9 ± 7.0 μm) × 1-2 μm width (av. ±S.D = 1.5 ± 0.4 μm) (n = 50), (L/W ratio = 14.6).

Based on morphological characters and pathogenicity tests, four representative isolates of *Ladiodiplodia* sp. (Fr-F01; FRK37,FRZ20 and FRZ27) and one of *Collethotrichum* sp.(SA9FSy), *Curvularia* sp. (BenFFSy), *Fusarium* sp. (1BRaA2) and *Diaporthe* sp. (NTA) were selected for molecular identification (Table 2).

**Table 2:** Characteristics of the isolates selected for molecular identification of fungi associated with mango Stem-end rot.

| Isolate | Species* | Length of<br>conidia (µm) | Width of<br>conidia (µm) | Average L/W<br>of conidia | GenBank Accession<br>numbers |  |
| --- | --- | --- | --- | --- | --- | --- |
|  |  |  |  |  | ITS | tef1-α |
| BenFFSy | *Curvularia pseudobrachyspora* | 17.2-26 | 9-11 | 2.7 | OR264470 | OR287046 |
| SA9FSy | *Colletotrichum gloeosporioides* | 15-18.0 | 5-6 | 3 | OR264469 | OR338285 |
| 1BRaA2 | *Fusarium mangiferae* | Macroconidia<br>(22.5-37.1) | 2-3.6 | 10.6 | OR264471 | OR287047 |
| NTA | *Diaporthe endophytica* | Beta conidia<br>(19-25) | 1-1.6 | 16.9 | OR264472 | OR287048 |
| Fr-F01 | *Lasiodiplodia theobromae* | 19.8-27.7 | 10.8-14.2 | 1.8 | OQ067836 | OQ269654 |
| Fr-K37 | *Lasiodiplodia euphorbicola* | 16.5-31.6 | 11.9-15.22 | 1.9 | OQ067837 | OQ269655 |
| Fr-Z20 | *Lasiodiplodia euphorbicola* | 20.7-26.4 | 10-14.3 | 1.9 | OQ067838 | OQ269656 |
| Fr-Z27 | *Lasiodiplodia caatinguensis* | 15.4-25.5 | 12.1-15.1 | 1.7 | OQ067839 | OQ269657 |
| *As determined by *tef1-α* sequence analysis |  |  |  |  |  |  |

The isolates were then subjected to multigene DNA sequence analysis and identified at the species level (Table 1). The multigene DNA sequence analysis based on ITS and *tef1*-α demonstrated that the SER sequences were similar to the reference sequences of *Lasiodiplodia theobromae* (FRF01), *L euphorbicola* (FRK37, FRZ20), *L. caatinguensis* (figure 4, supplementary figure 1), *Colletotrichum gloeosporioides* (SA9FSy) (figure 5; supplementary figure 2), *Curvularia pseudobrachyspora* (BenFFSy) (figure 6; supplementary figure 3), *Fusarium mangiferae* (1BRaA2) (figure 7; supplementary figure 4) and *Diaporthe endophytica* (NTA) (figure 8; supplementary figure 5).

**Figure 4:**
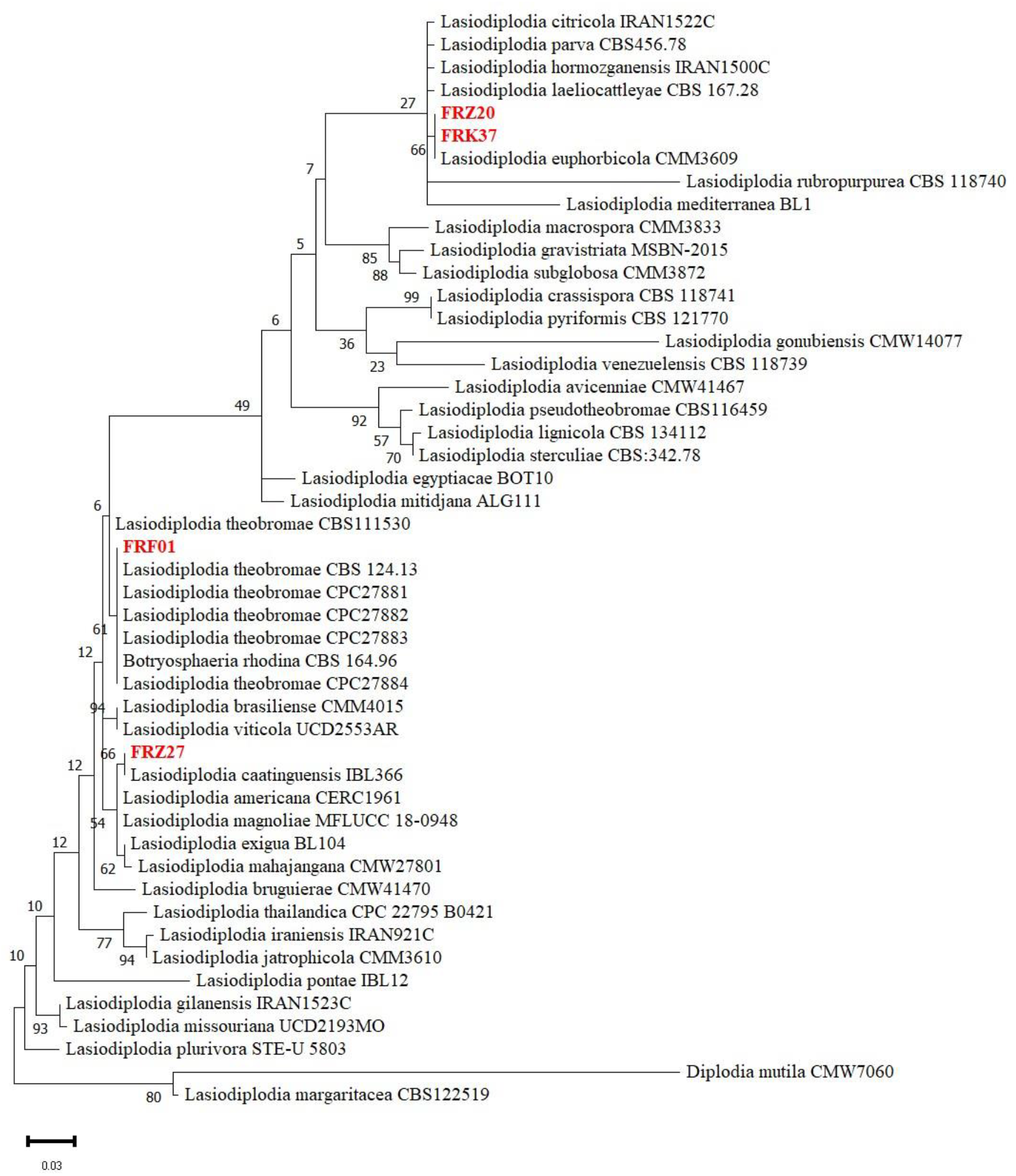
Maximum likelihood tree of *Lasiodiplodia* species isolated from mango SER based on *tef1*-α sequences dataset using Kimura 2-parameter model with 1000 bootstrap replications. The species name is followed by the strain accession numbers, and the isolates from the current study are in red. The bootstrap support values above 50% are given at the node. There were a total of 296 positions in the final dataset. *Diplodia mutila* (CMW7060) is used as the outgroup taxon.

**Figure 5:**
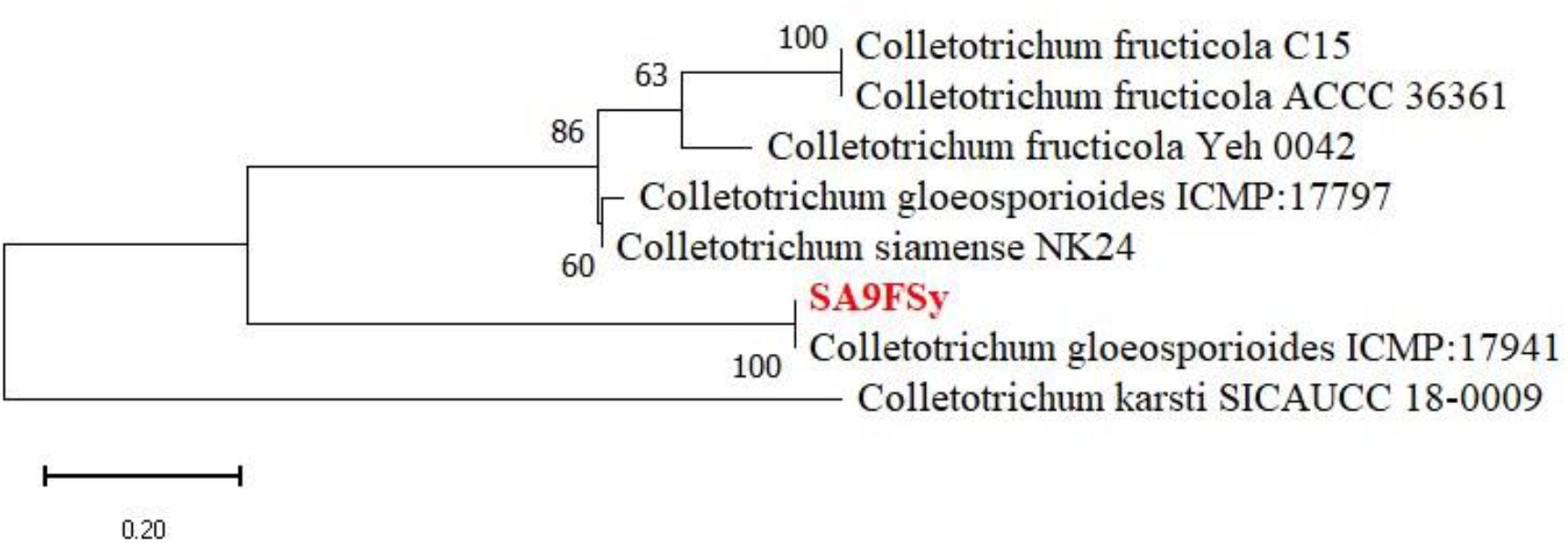
Maximum likelihood tree of *Colletotrichum* species based on the *tef1*-α sequences dataset using Kimura 2-parameter model with 1000 bootstrap replications. The species name is followed by the sequence accession numbers, and the isolate from the current study are in red. The bootstrap support values above 50% are given at the nodes. There were a total of 367 positions in the final dataset. *Colletotrichum karsti* (SICAUCC) is used as the outgroup taxon.

**Figure 6:**
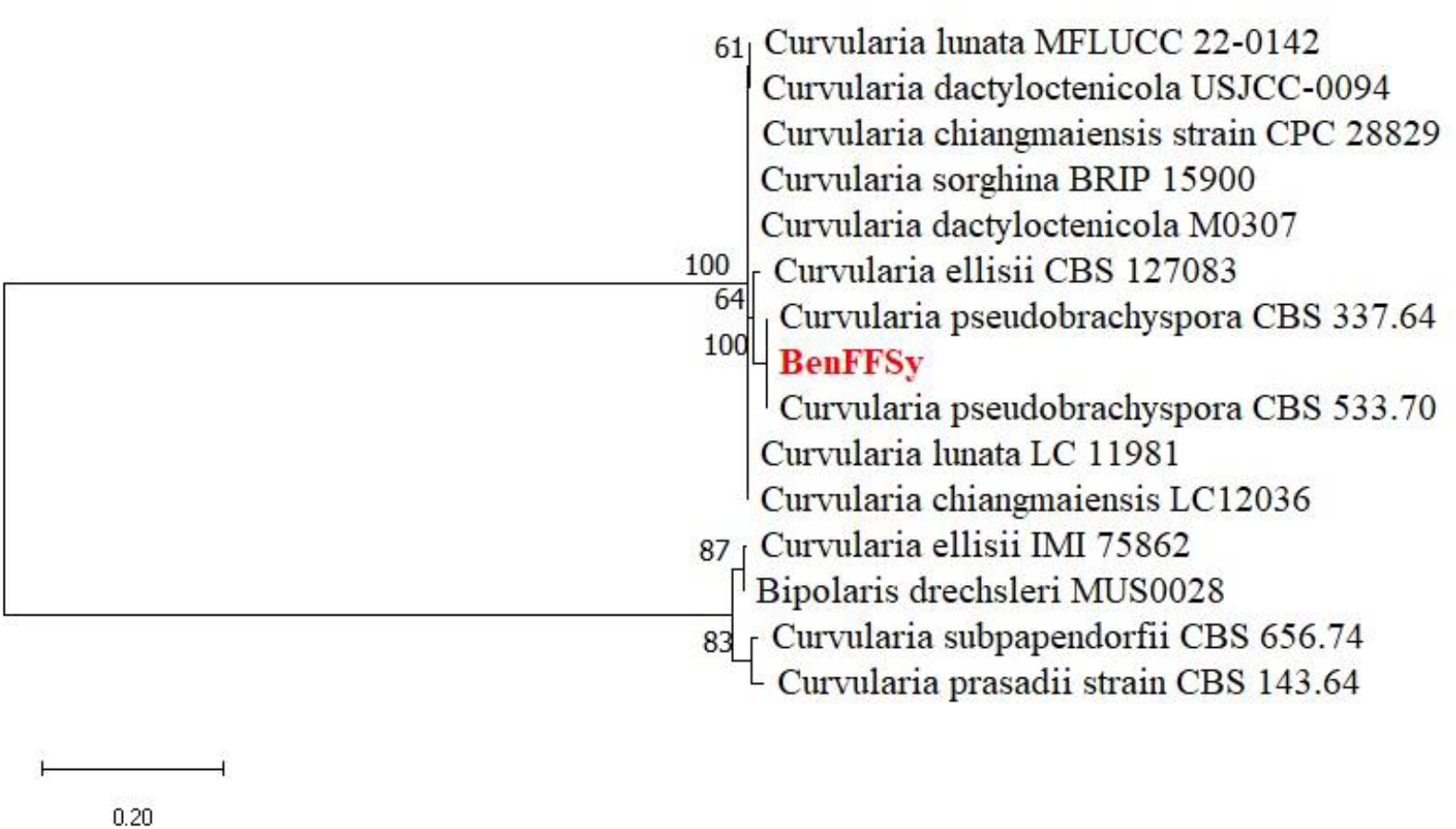
Maximum likelihood tree of *Curvularia* species based on the *tef1*-α sequences dataset using Kimura 2-parameter model. With 1000 bootstrap replications. The species name is followed by the sequence accession numbers, and the strain from the current study are in red. The bootstrap support values above 50% are given at the node. There were a total of 877 positions in the final dataset. *Bipolaris drechsheri* (MUSOO28) is used as the outgroup taxon.

**Figure 7:**
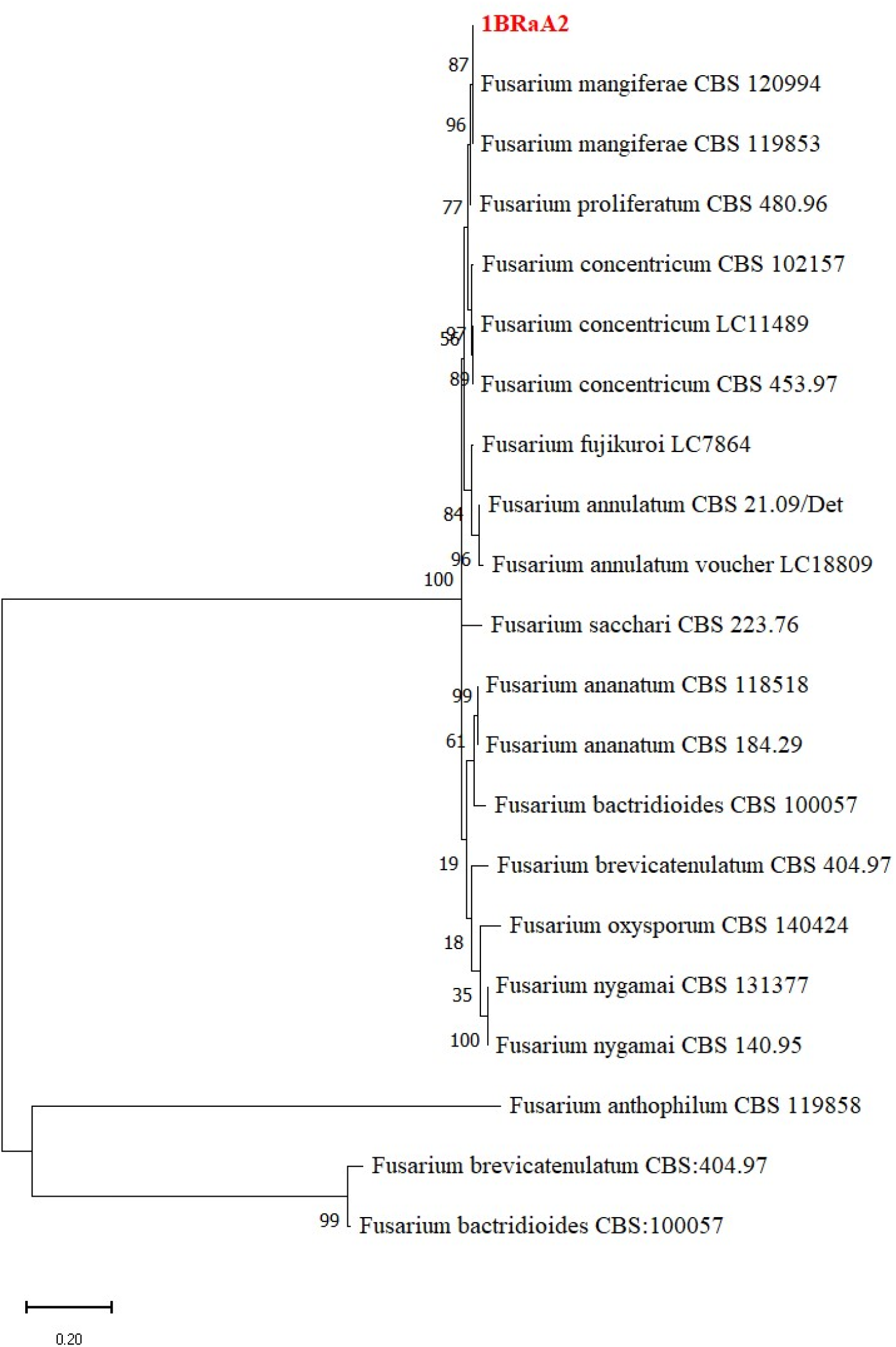
Maximum likelihood tree of *Fusarium* species based on the *tef1*-α sequences dataset using Jukes-Cantor modele with 1000 bootstrap replications. The species name is followed by the sequence accession number, and the isolate from the current study is in red. There were a total of 622 positions in the final datase. *Fusarium bactridioides* (CBS 100057) is used as the outgroup taxon.

**Figure 8:**
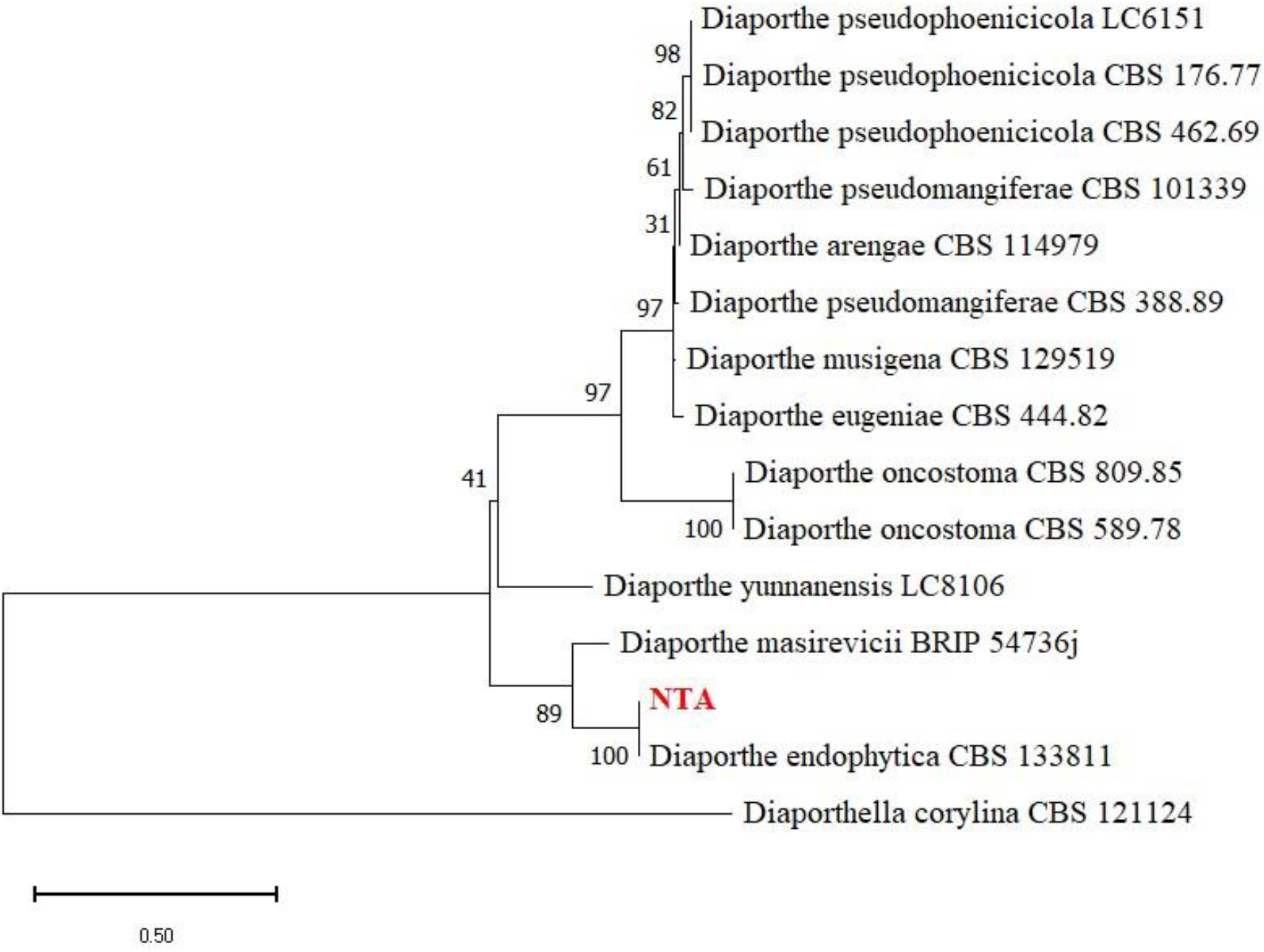
Maximum likelihood tree of *Diaporthe* species based on the *tef1*-α sequences dataset using Hasegawa-Kishino-Yano model with 1000 bootstrap replications. The species name is followed by the sequence accession numbers, and the NTA strain from the current study is in red. The bootstrap support values above 50 % are given at the nodes. There were a total of 374 positions in the final dataset. *Diaporthe corylina* (CBS 121124) is used as the outgroup taxon.

### 3.4 Pathogenicity tests

The eight SER isolates inoculated on mango fruits induced peel discoloration during the incubation period. However, only *Lasiodiplodia* (FrF01, FrK37, FrZ20, FrZ27) and *Diaporthe* (NTA) isolates induced characteristics SER symptoms on the artificially inoculated fruits, similar to those observed on naturally infected fruits. SER symptoms included a water-soaked, dark, discoloured area near the fruit stem, which could range from dark brown to purplish-black (Figure 4). These symptoms usually appeared on the ripe or ripening fruit peel at the stem-end region. At first, a dark-brown to black ring appears around the pedicel, which expanded as the fruit ripened, covering the upper one-third of the fruit peel (Figure 4a-e). As the infection progressed, the affected areas quickly darkened and merged, resulting in complete fruit rot 10 days after inoculation (DAI) (Figure 4f). The fungi were reisolated on PDA, and the colony morphology and conidia produced in cultures were identical to the isolates of each species used originally for fruit inoculation.

**Figure 4:**
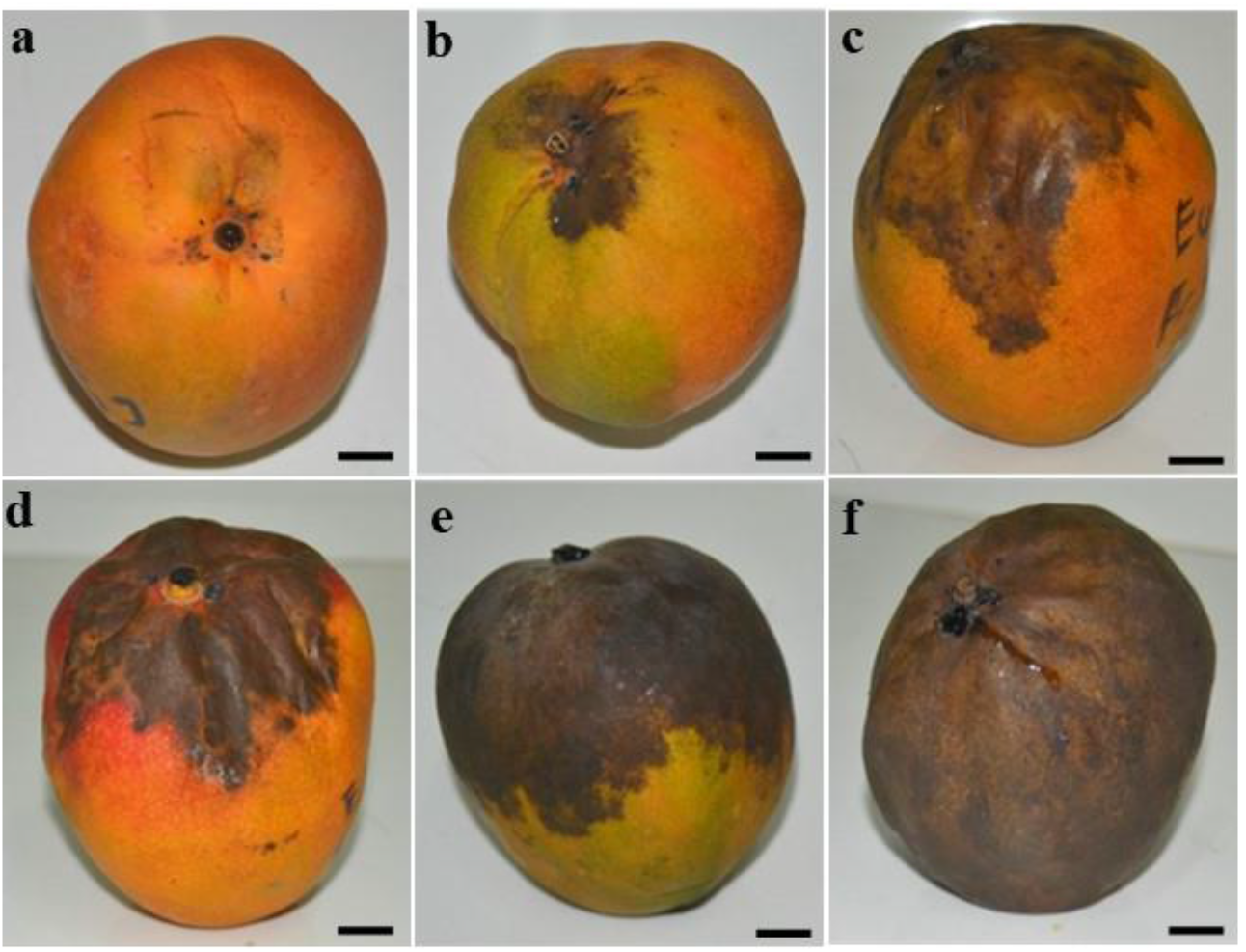
Evolution of mango Stem-end rot (SER) symptoms on mango cv Kent fruits. **a**= First day of symptoms apparition:discoloration limited at the stem-end at 3 days after inoculation; **b**= 10% discoloration of the fruit surface area initiated by stem-end rot at 4 days after inoculation; **c=** 11-30% discoloration of the fruit surface area initiated by stem-end rot at 7 days after inoculation; **d**= 31-50% discoloration of the fruit surface area initiated by stem-end rot at 8 days after inoculation; **e**= more than 51% discoloration of the fruit surface area initiated by stem-end rot at 9 days after inoculation; **f**= fruit completely rotted (100 %) after 10 days after inoculation. Scales Bars = 2 cm.

The lesion diameter of the fruits showing SER symptoms varied from 5 to 29 cm at 10 DAI (figure 5). *Lasiodiplodia* species induced the highest diameters (more than 28,5 cm), while the lesion diameters induced by *Diaporthe endophytica* were the lowest (5 cm) (P <.0.05). (Figure 5). There was no significant difference between the lesion diameters induced by different *Lasiodiplodia* isolates.

**Figure 5:**
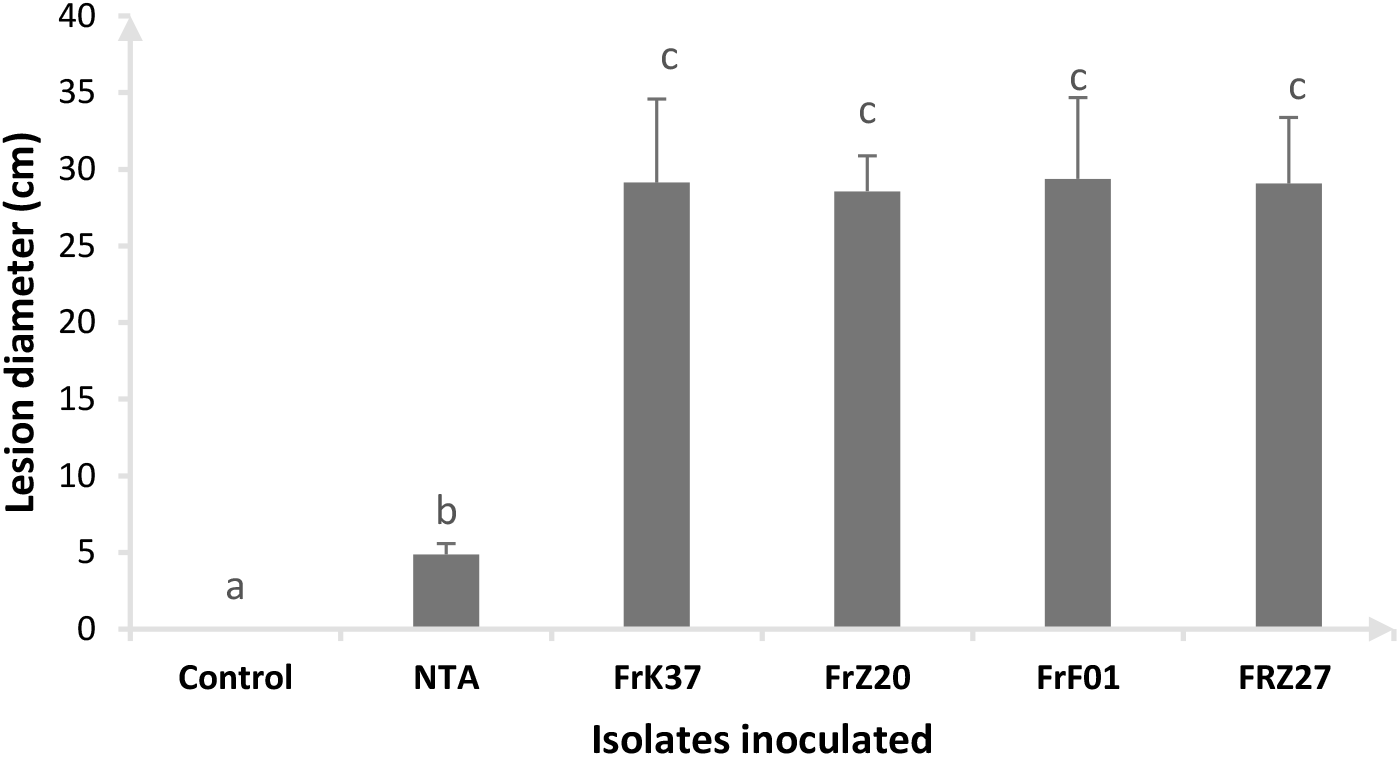
Mango fruit lesion diameter at 10 days after inoculation with SER representatives isolates. NTA= *Diaporthe endophytica* FrK37, FrZ20 = *Lasiodiplodia euphorbicola*; FrF01= *L. theobromae*; FrZ27= *L. caatinguensis*. Values presented are mean ±SD (*n=15*). Values with different letters show a significant difference by ANOVA (p≤0.05), LSD test at p=0.05.

## 4 Discussion

Mango SER is a significant threat to all mango sector actors, notably farmers and export industries. Investigating disease epidemiology is imperative to control SER and limit mango quality losses. The current study evaluated mango SER intensity in the main production area in North West Côte d’Ivoire, determined the correlation between incidence and severity and preharvest climate parameters, and identified the SER causing fungi.

Symptoms of SER were characterized by the development of lesions that initially appeared as small, dark-brown to black areas located in the peel around the base of the fruit stem end. These lesions then progressed into a soft, watery rot that spread outwards from the peduncle, resulting in complete fruit rot after 10 days, as described by Galsurker et al. (2020) in Pakistan and Honger and William (2015) in Ghana. SER symptoms differ from anthracnose disease, which are characterized by black spots (Alvindia and Acda, 2015a; Honger and William, 2015).

Surveys conducted in 2020 and 2021in different production departments indicated that SER is prevalent in the North West agroecological zone of the country. Depending on orchards, SER incidence varied between 10 to 30 % while severity was 5 to 20 % and both variables were strongly correlated (Figure 1). Some previous studies had reported similar values, notably those conducted in Pakistan by Syed et al. (2017) and Israel by Feygenberg et al. (2021). In addition, despite the SER incidence and severity varied between departments, no significant difference was found between them for both variables, suggesting that the geographical location may not significantly impact mango SER in this agroecological zone of Côte d’Ivoire. This result could be explained because the same cultural practices and agroclimatic conditions are prevailing within the selected orchards, as proposed by Syed et al. (2017) in Pakistan.

Storage temperature and relative humidity effect on mango SER incidence were reported by many authors worldwide (Alam et al., 2017b; Alvindia and Acda, 2015a; Diskin et al., 2017). However, there is limited data on the preharvest weather parameters effect on mango SER. Our study analyzed the correlation between climate parameters recorded during a six-months preharvest period and SER incidence and severity in orchards. We found a low positive correlation between the mean temperature (T2M) and the minimum temperature (T2M_MIN) with SER incidence (r = 0.36 and r = 0.26, respectively). The current results agree with many authors demonstrating that environmental conditions are relevant variables affecting pathogen infection (Riquelme-Toledo et al., 2020). Indeed, Honger and William (2015) identified that the district in Ghana with consistently high temperatures throughout the year had the highest mango SER. Based on our statistical analysis, the absence of correlation between mango SER incidence and severity with precipitation and relative humidity suggest that other factors, such as farmers’ cultural practices and soil type, may better explain the intensity of the disease in these regions of Côte d’Ivoire.

The present study identified several fungal genera associated to mango SER, including *Lasiodiplodia* sp*., Curvularia* sp., *Fusarium* sp., *Colletotrichum* sp., *Diaporthe* sp. Among them, isolates belonging to the *Lasiodiplodia* genus were the most frequently isolated, with a 74% isolation rate. Assignment to species based on multigene phylogenetic analysis of ITS and *tef1-α* sequences may not be accurate for those genera where cryptic species have been recently denominated (Alves et al., 2008; Dissayanake et al., 2016). The species concept for filamentous fungi is still largely discussed and debated (Stengel et al., 2022). In this study, sequences obtained for SER isolates were identical to sequences deposited in GenBank for type isolates from seven species, namely *Lasiodiplodia theobromae, L. caatinguensis*, *L. euphorbicola, Colletotrichum gloeosporioides*, *Fusarium mangiferae*, *Curvularia pseudobrachyspora* and *Diaporthe endophytica*. However, these sequences also matched other closely related fungal species with the same identity level. Therefore without investigating further the genetic diversity present in a given species, it is questioned whether some closely related species may not be erroneously assigned to different species based on a few genetic markers, such as ITS and *tef1-α* sequences.

All the SER isolates tested in this study were pathogenic to fruits. However, only *Lasiodiplodia* species and *Diaporthe endophytica* isolates exhibited true SER symptoms, and *Lasiodiplodia* species were the most aggressive. *Lasiodiplodia* species might have a high rate of mango inflorescence colonization which could explain the highest isolation percentage. Furthermore, according to Coelho et al. (2020) and Ji et al. (2017), *Lasiodiplodia* species infect many plant species, including fruit trees, grapevines, ornamental plants, and forest trees. Our results are in agreement with those of Karunanayake and Adikaram (2020) showing that *Lasiodiplodia* species are the major pathogens causing SER disease in mango, mainly in the dry and warmer regions of the world. However, several studies have reported that it is responsible for several other destructive diseases, such as canker, dieback and gummosis (Quimbita-Reyes et al., 2023; Coelho et al., 2020; Netto et al., 2017).

As in the present study, the fungal genera *Colletotrichum* sp., *Curvularia* sp., *Diaporthe* sp., and *Fusarium* sp. were reported to be associated with mango SER in several previous studies (Ajitomi et al., 2020; Alam et al., 2017; Guarnaccia, 2016; Honger and William, 2015). However, *Colletotrichum gloeosporioides* is mainly known as the causal agent of mango anthracnose, which is characterized by small, circular, sunken spots that appear on the mango fruit surface (Dembélé et al., 2020; N’Guettia et al., 2013). Nevertheless, the symptoms are completely different from those of SER and should not be confused with it (Honger and William, 2015). Recently, Adikaram et al. (2022) discovered *Curvularia* species causing stem-end browning symptoms (SEB) in mango, which are distinct from SER symptoms. About *Diaporthe* sp (syn *Phomopsis*), the main species reported on mango SER were *Diaporthe eugeniae*, *D. pascoei*, *D. perseae*, and *D. ueckerae* (Ajitomi et al., 2020; Lim et al., 2019). Our isolate was similar to *Diaporthe endophytica*, which has been isolated on SEB in ripe mango by Adikaram et al. (2022a). *Fusarium mangiferae* were identified on SER symptoms, while Alvindia and Acda (2015) reported *F. verticillioides*. *F. mangiferae* was also reported as responsible for mango malformation in Sri Lanka (Kausar et al., 2021) and SEB disease in ripe mango (Adikaram et al., 2022a).

The presence of all these isolates concurrently isolated on SER symptoms could be explained by the fact that SER symptoms expanded and masked the lesions of the other fungal, as mentioned in the study conducted by Honger and William (2015).

## Conclusion

The present study focused on mango SER in Côte d’Ivoire, one of the most important postharvest disease in mango production areas worldwide. This study showed that mango SER is prevalent in Côte d’Ivoire, with an incidence rate of up to 30 %. In addition, meteorological factors such as preharvest temperature are among the parameters that can influence this disease. Therefore, environmental factors could be included in the panel of control methods for SER. Furthermore, seven species of fungal pathogens were identified, with Botryosphaeriaceae (*Lasiodiplodia* spp) the most isolated family. By filling the knowledge gap on the epidemiology of the SER disease of mango in Côte d’Ivoire, our results establish a solid basis to guide future research on mango postharvest diseases. Therefore, a broader study and evaluation of mango plantations should be undertaken in future studies to better understand the relationship between overall mango infection and postharvest development of SER.

## ACCESSION NUMBERS

Sequence data from this article listed in table 2 can be found in the GenBank data libraries under the different accession numbers mentioned for ITS and tef1-α genes.

## FUNDING

This research was funded by the Partnership for Skills in Applied Sciences, Engineering, and Technology—Regional Scholarship and Innovation Fund (PASET-RSIF), the French Research Institute for Development (IRD, France), and the Fondation Louis Omer DECUGIS (OMER-DECUGIS & CIE - 94538 RUNGIS, FRANCE).

## ACKNOWLEDGEMENTS

The first author acknowledges the support from the Société Internationale d’ Importation (SIIM, France) and the Société de Diverses Prestations et d’Exportations (Sodipex SARL, Côte d’Ivoire) for their technical support. We also thank mango producers for the facilities they provided us during the survey.

## AUTHOR CONTRIBUTIONS

Yéfoungnigui S. YEO: Conceptualization; Methodology; Resources; Formal analysis; Roles/Writing - original draft

Daouda KONE: Conceptualization; Funding acquisition. Project administration

Yassogui KONE: Methodology

Dio D. DEMBELE: Review & editing

Elisee L D G AMARI: Roles/Writing - review & editing

Jean-Yves REY: Funding acquisition; Writing - review & editing

Emerson M. DEL PONTE : Methodology; Formal analysis; Roles/Writing - review & editing

Diana FERNANDEZ: Conceptualization; Funding acquisition; Methodology; Formal analysis; Roles/Writing - review & editing

## CONFLICT OF INTEREST STATEMENT

The authors declare no conflict of interest.

## Supplementary figures

**Supplemental Figure 1:**
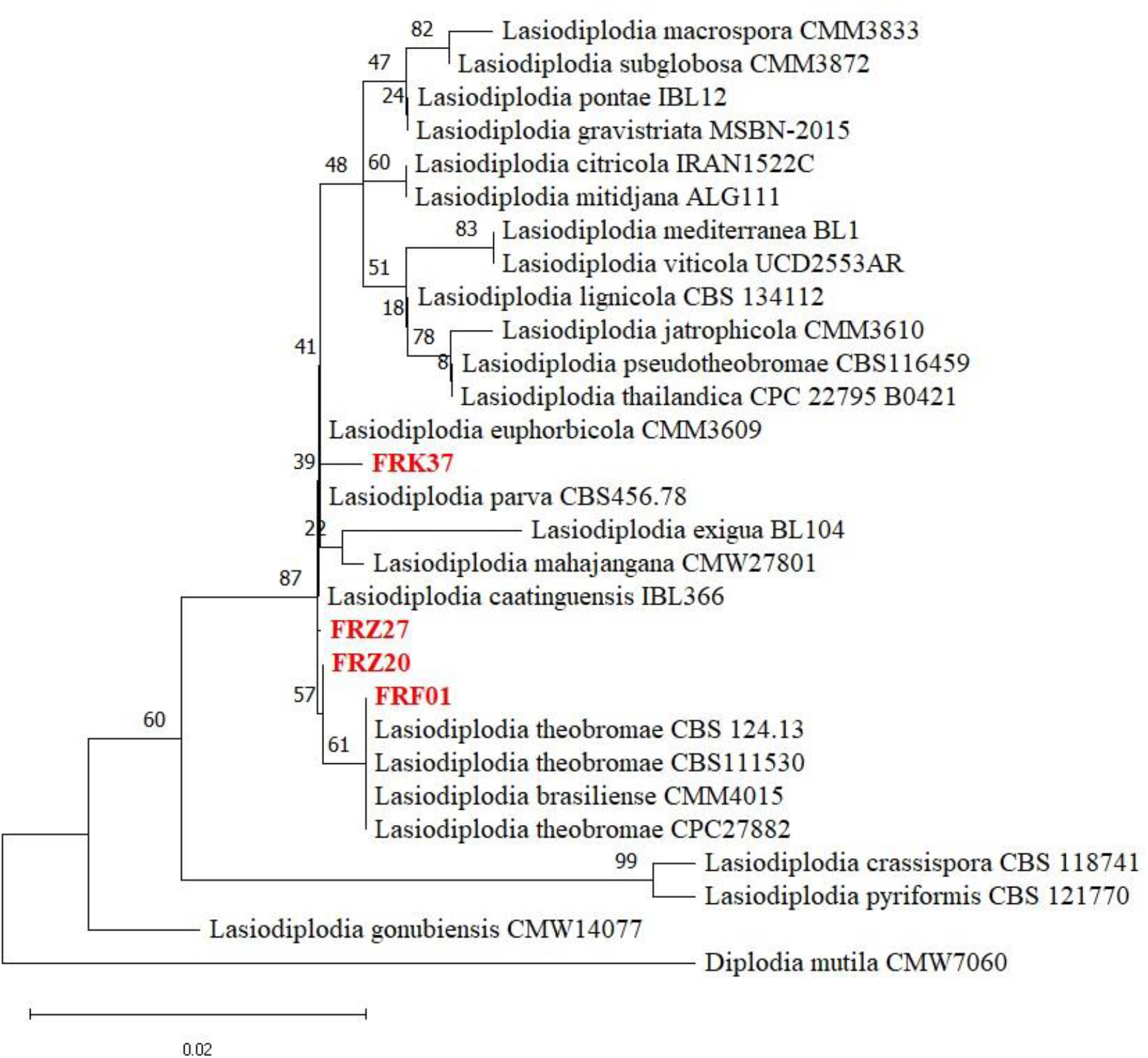
Phylogram generated by Maximum likelihood based on ITS sequences data of 4 *Lasiodiplodia* spp. isolated on mango SER and 25 *Lasiodiplodia* references species from the NCBI database. The species name is followed by the strain accession numbers, and the strain from the current study are in red. The bootstrap support values above 50% are given at the nodes. The evolutionary history was inferred by using the Maximum Likelihood method and Kimura 2-parameter model [1]. The tree with the highest log likelihood (−884.45) is shown. Initial tree(s) for the heuristic search were obtained automatically by applying the Maximum Parsimony method. There were a total of 499 positions in the final dataset. *Diplodia mutila* (CMW7060) is used as the outgroup taxon.

**Supplemental Figure 2:**
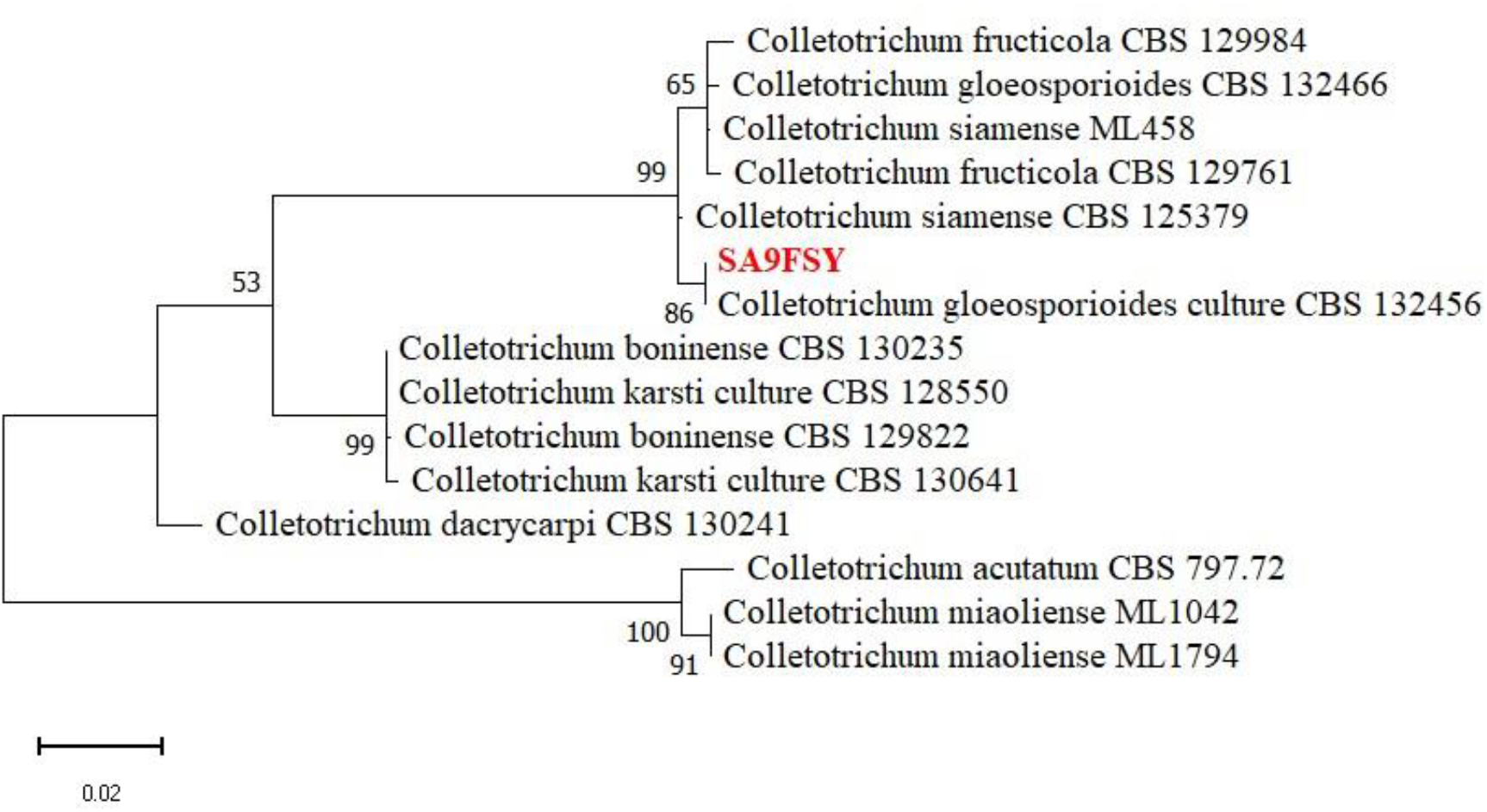
Phylogram generated by Maximum likelihood based on the ITS sequences data of the isolate SA9FSY isolated from mango SER and 14 *Colletotrichum* references species from the NCBI database. The species name is followed by the sequence accession numbers, and the strain from the current study are in red. The bootstrap support values above 50% are given at the nodes. The evolutionary history was inferred by using the Maximum Likelihood method and Kimura 2-parameter model [1]. The tree with the highest log likelihood (−1387.81) is shown. Initial tree(s) for the heuristic search were obtained automatically by applying the Maximum Parsimony methods. There were a total of 585 positions in the final dataset. *Colletotrichum miaoliense* (ML1794) is used as the outgroup taxon.

**Supplemental Figure 3:**
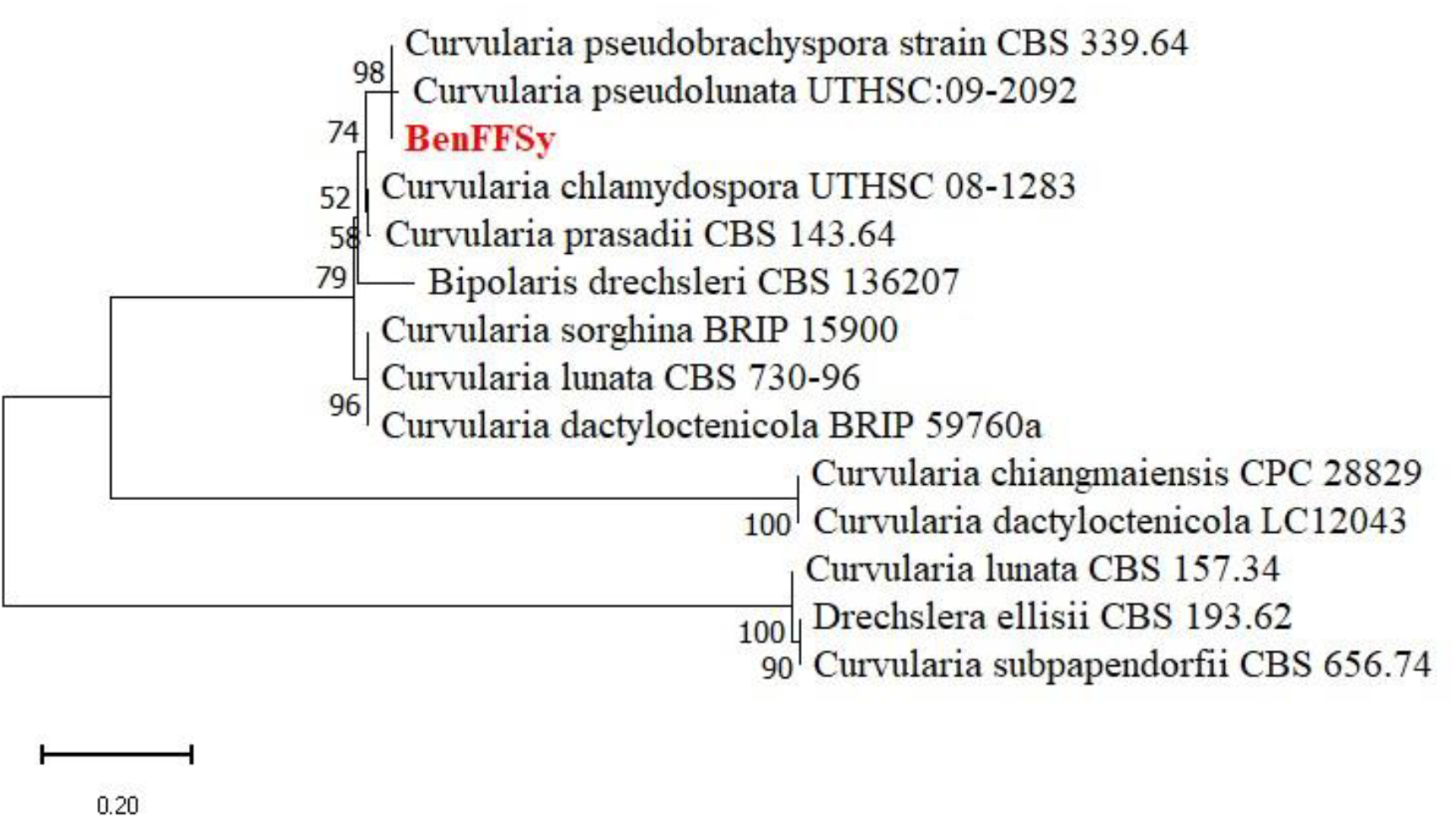
Phylogram generated by Maximum likelihood based on the ITS sequences data of the isolate BenFFSy isolated from mango SER and 13 *Curvularia* references species from the NCBI database. The species name is followed by the sequence accession numbers, and the strain from the current study are in red. The bootstrap support values above 50% are given at the nodes.The evolutionary history was inferred by using the Maximum Likelihood method and Kimura 2-parameter model [1]. The tree with the highest log likelihood (−2343.11) is shown. There were a total of 635 positions in the final dataset. *Curvularia lunata* (CBS 557.34) is used as the outgroup taxon.

**Supplemental Figure 4:**
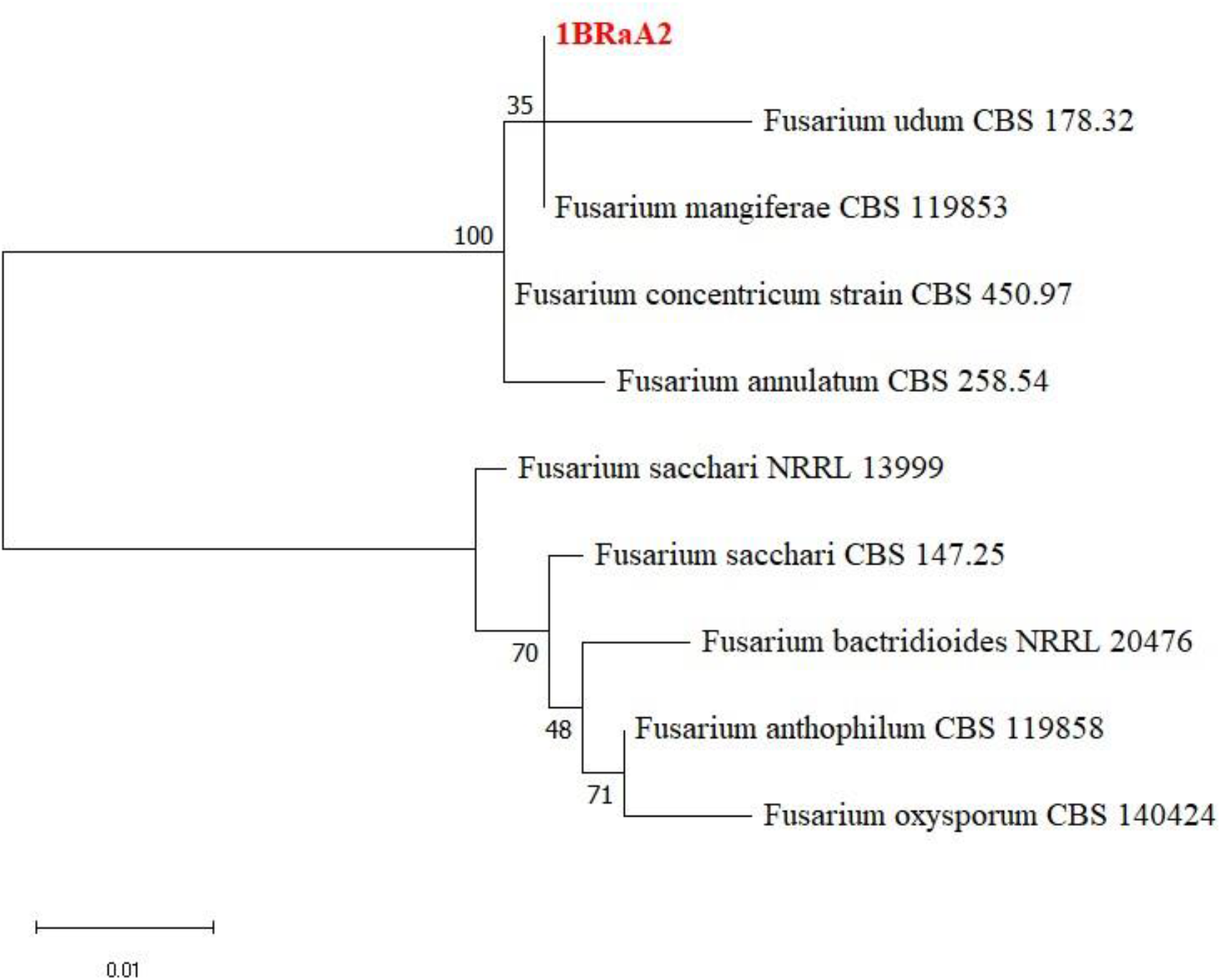
Phylogram generated by Maximum likelihood based on the ITS sequences data of the isolate 1BRaA2 isolated from mango SER and 9 *Fusarium* references species from the NCBI database. The species name is followed by the sequence accession number, and the strain from the current study is in red. The bootstrap support values above 50% are given at the nodes. The evolutionary history was inferred by using the Maximum Likelihood method and Jukes-Cantor modele. The tree with the highest log likelihood (−1262.13) is shown. There were a total of 749 positions in the final datase. *Fusarium oxysporum* (CBS 140424) is used as the outgroup taxon.

**Supplementary Figure 5:**
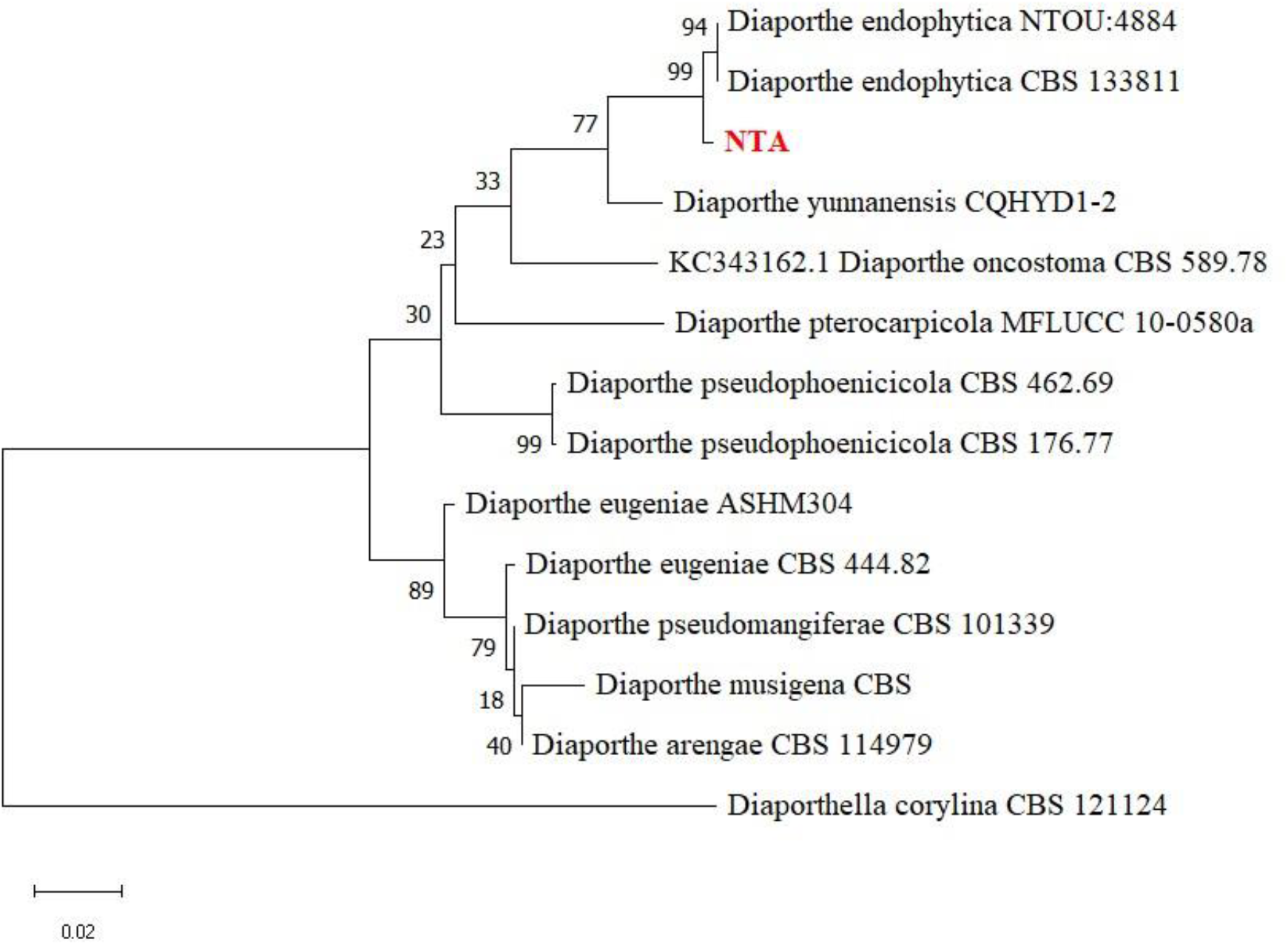
Phylogram generated by Maximum likelihood based on the ITS sequences data of NTA isolate isolated from mango SER and 13 *Diaporthe* references species from the NCBI database. The species name is followed by the sequence accession numbers, and the NTA strain from the current study is in red. The bootstrap support values above 50 % are given at the nodes. The evolutionary history was inferred by using the Maximum Likelihood method and Kimura 2-parameter model. The tree with the highest log likelihood (−1772.67) is shown. Initial tree(s) for the heuristic search were obtained automatically by applying the Maximum Parsimony method. There were a total of 633 positions in the final dataset. *Diaporthe corylina* (CBS 121124) is used as the outgroup taxon.

## Notes

### Competing Interest Statement

The authors have declared no competing interest.

